# Brief Communication: Differential degradation of exogenous DNA in microalgal cell lysates

**DOI:** 10.64898/2026.09.23.753703

**Authors:** Ekta Kumari, Nanakow Baiden, John Verruto, Oscar Marsden, John Waite, Ian Hu

**Affiliations:** PHYCOBLOOM LTD. Unit 130 Westbourne Studios, 242 Acklam Road, London, England, W10 5JJ

**Keywords:** *Chlorella sorokiniana*, genetic transformation, intracellular nuclease

## Abstract

*Chlorella sorokiniana* is an industrially important microalga, but its genetic manipulation remains limited by poor transformation efficiency. Although several barriers have been proposed, the role of intracellular nuclease activity has received little attention. Here, we compared the degradation of exogenous double-stranded DNA in whole-cell lysates from transformation-recalcitrant (*Chlorella sorokiniana* and *Tetradesmus obliquus*) and transformation-proficient (*Phaeodactylum tricornutum, Microchloropsis gaditana*, and *Nannochloropsis oceanica*) microalgae. *C. sorokiniana* rapidly degraded both linear and plasmid DNA, while *T. obliquus* exhibited intermediate activity. In contrast, no detectable degradation was observed in the transformation-proficient species. DNA degradation in *C. sorokiniana* occurred within minutes, and nuclease activity in both recalcitrant species was inhibited by EDTA and Zn^2+^, consistent with a metal ion-dependent endonuclease. These findings suggest that intracellular nuclease activity may represent an underexplored barrier to exogenous DNA stability during transformation and provide a potential target for improving genetic transformation of recalcitrant microalgae.

## 1 Introduction

*Chlorella sorokiniana* is one of the most industrially promising microalgae due to its rapid growth, high biomass productivity, and exceptional thermotolerance. It grows optimally at 35–40 °C, with greater tolerance to even more elevated temperatures compared with many *Chlorella* species, including *C*. vulgaris, making it well suited for large-scale outdoor cultivation in diverse environments ^1–3^. In addition to its robust growth, *C. sorokiniana* can accumulate substantial amounts of lipids, making it an attractive species for sustainable biofuel production and other biotechnological applications ^2,4^. Realising the full industrial potential of this species, as well as advancing our understanding of its biology, requires efficient and reliable genetic manipulation.

Despite its industrial potential, genetic engineering of *C. sorokiniana* remains challenging due to low, strain-dependent transformation efficiency and the limited availability of robust genetic tools ^5–8^. Transformation outcomes vary considerably between strains and laboratories, limiting reproducibility and broader application. In contrast, microalgae such as *Phaeodactylum tricornutum* are genetically tractable models with established molecular toolkits that support stable transgene expression, genome editing, and synthetic biology applications ^9,10^. Ongoing efforts to improve transformation in *C. sorokiniana* highlight the need for further advances. Several factors have been proposed to limit transformation, including the cell wall, codon usage bias, and transgene silencing ^5,8^. However, one potential barrier that has received little attention is the host defence system, particularly intracellular nuclease that may degrade incoming foreign DNA before stable integration can occur.

Intracellular nucleases degrade exogenous nucleic acids and are recognized as an important factor that can limit gene delivery efficiency in diverse biological systems ^11,12^. In mammalian cells, cytoplasmic nucleases degrade microinjected plasmid DNA with a half-life of 50–90 minutes, constituting a major barrier to non-viral gene delivery ^12^ . Nuclease-mediated degradation of exogenous DNA has been proposed as a contributing factor influencing transformation outcomes in some microalgae. In *Dunaliella tertiolecta*, introduced plasmid DNA is degraded following electroporation, while in *Desmodesmus armatus*, introduced DNA undergoes extensive truncation and rearrangement during integration, consistent with nuclease-mediated degradation of foreign DNA ^13,14^ . Ca^2+^-dependent intracellular nucleases have been characterized in *Chlamydomonas reinhardtii*, showing preference for single-stranded DNA ^15–17^. A similar preference for denatured DNA has been reported in a strain historically identified as *Chlorella pyrenoidosa* and now classified as *Coelastrella vacuolata*, with substantially lower nuclease activity observed toward native DNA ^18^.

To our knowledge, intracellular nuclease activity has not been directly compared across microalgal species with contrasting transformation tractability, nor has its potential contribution to exogenous DNA stability been systematically investigated. Here, we compare double-stranded DNA degradation in whole-cell lysates from transformation-recalcitrant species (*Chlorella sorokiniana* and *Tetradesmus obliquus*) and genetically tractable reference species (*Microchloropsis gaditana, Nannochloropsis oceanica*, and *Phaeodactylum tricornutum*) using plasmid and linear DNA substrates. By relating nuclease activity to transformation tractability, we assess whether intracellular DNA degradation represents an underexplored barrier to successful genetic transformation in industrially relevant microalgae.

## 2 Materials and Methods

### 2.1 Algal strains and culture conditions

The algal strains used in this study were *Chlorella sorokiniana* (SAG 211-32), *Tetradesmus obliquus* (CCAP 276/7 and CCAP 258/164), *Phaeodactylum tricornutum* (CCMP 632), *Microchloropsis gaditana* (CCAP 849/5), and *Nannochloropsis oceanica* (CCAP 849/8). The freshwater species *C. sorokiniana* and *T. obliquus* were cultured in Tris-acetate-phosphate (TAP) medium, while the marine species *P. tricornutum, M. gaditana*, and *N. oceanica* were cultured in artificial seawater (ASW) medium. Freshwater species were maintained at 25 °C and marine species at 18 °C. All cultures were grown at a light intensity of 150 μmol photons m^−2^ s^−1^ under a 16 h light/8 h dark photoperiod.

### 2.2 Nuclease assay

Nuclease activity was assessed as previously described, with minor modifications^13^. Cultures were harvested during the logarithmic growth phase. Cells were collected by centrifugation and washed twice with 10 mM Tris-HCl (pH 8.0) containing 50 mM NaCl at 4 °C. Cell pellets were maintained on ice throughout the extraction procedure. Cells were resuspended in the same buffer supplemented with 5% glycerol, 0.1% Triton X-100, and cOmplete™ protease inhibitor (Roche, USA; Cat. No. 04693159001). Cell suspensions were lysed using glass beads (Merck; Cat. No. Z763721-50EA).

Following lysis, the cell suspension was centrifuged at 13,000 × *g* for 15 min at 4 °C. The resulting supernatant was collected, and protein concentration was determined using the Pierce BCA Protein Assay Kit (Thermo Fisher Scientific; Cat. No. 23225). For the nuclease assay, 10 μg of total protein was incubated with 500 ng DNA (Plasmid DNA or Lambda DNA) in 1× Tango buffer (Thermo Scientific; Cat. No. BY5), and the reaction volume was adjusted to 40 µL with molecular-grade water. Reaction mixtures were incubated at 30 °C for different time periods, as indicated in the Results and Discussion section. Following incubation, samples were analysed by electrophoresis on a 1% agarose gel prepared in Tris-borate EDTA (TBE) buffer to assess DNA degradation.

For the nuclease inhibition assays, EDTA and ZnSO_4_·7H_2_O were added to the reaction mixtures at final concentrations of 10 mM and 5 mM, respectively. The reactions were incubated at 30 °C for the time periods indicated in the Results and Discussion section. Nuclease activity was assessed by agarose gel electrophoresis as described above.

## 3 Results and Discussion

### 3.1 Species-dependent nuclease activity in algal cell lysates

To investigate whether species-dependent nuclease activity may represent a barrier to exogenous DNA stability during transformation, we compared exogenous DNA degradation in whole-cell lysates from algal species with contrasting transformation tractability. *P. tricornutum, M. gaditana*, and *N. oceanica*, which possess established transformation protocols and advanced genetic engineering toolkits, were selected as transformation-proficient reference species ^19– 22^. In contrast, *C. sorokiniana* and two *T. obliquus strains* (hereafter referred to as TO/7 and TO/164), which remain more challenging for stable genetic transformation and have less developed genetic engineering resources, were included to assess whether nuclease activity may contribute to reduced exogenous DNA stability ^23–25^.

To determine whether algal cell lysates contain nucleases capable of degrading exogenous DNA, an in vitro nuclease assay was performed using two DNA substrates with different topologies: linear λ DNA and circular plasmid DNA, following 24 h incubation. The EcoRI positive control confirmed that the assay conditions permitted DNA cleavage, generating the expected restriction fragments, while the no-lysate negative control retained intact DNA, confirming that substrate degradation was dependent on the presence of algal lysate. As shown in Figure 1, *C. sorokiniana* exhibited the highest detectable nuclease activity, with no detectable intact λ DNA (Figure 1A) or plasmid DNA (Figure 1B) remaining after incubation, resulting in a smeared DNA profile. Both *T. obliquus* strains (TO/7 and TO/164) also displayed nuclease activity, although less pronounced than in *C. sorokiniana*. λ DNA was extensively degraded with visible smearing, whereas plasmid DNA showed partial degradation, with reduced intensity of the upper band, increased intensity of the lower band, and visible smearing at 24 h incubation. In contrast, lysates from *P. tricornutum, N. oceanica*, and *M. gaditana* showed no detectable degradation of either substrate, indicating no measurable nuclease activity under the assay conditions.

**Figure 1.**
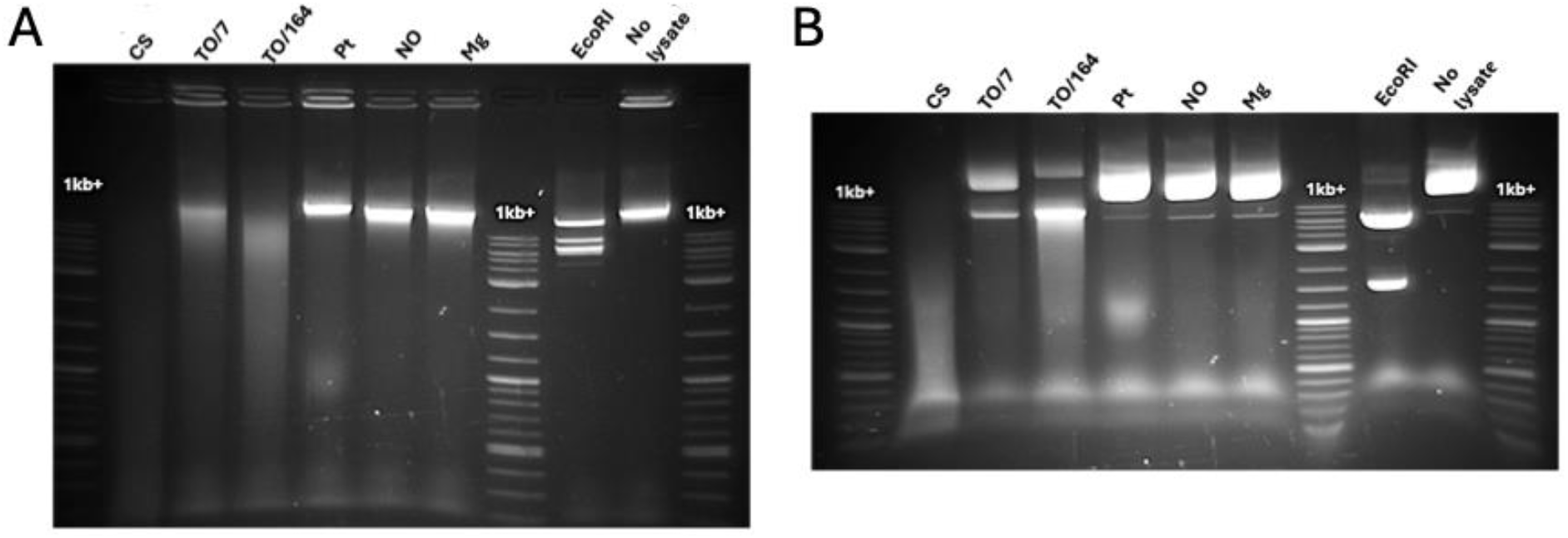
Agarose gel analysis of DNA degradation following incubation of DNA substrates with algal cell lysates. (A) Lambda DNA incubated with lysates for 24 h. (B) Plasmid DNA incubated with lysates for 24 h. The extent of DNA degradation varies among the tested algal species, revealing species-specific nuclease activity. Equal amounts of total protein from each lysate were used for comparison. Lanes are labelled as follows: CS, *Chlorella sorokiniana* (SAG 211-32/IBVF); TO/7, *Tetradesmus obliquus* (CCAP 276/7); TO/164, *Tetradesmus obliquus* (CCAP 258/164); Pt, *Phaeodactylum tricornutum* (CCMP 632); NO, *Nannochloropsis oceanica* (CCAP 849/8); Mg, *Microchloropsis gaditana* (CCAP 849/5). 1 kb+, 1 kb Plus DNA Ladder (New England Biolabs).

Collectively, these results reveal a gradient of nuclease activity that tracks with transformation tractability — highest in *C. sorokiniana*, intermediate in both *T. obliquus* strains, and undetectable in the three transformable reference species — suggesting that nuclease-mediated degradation of exogenous DNA may reduce DNA persistence following delivery. While transformation has been reported in *C. sorokiniana* and *T. obliquus*, these species remain considerably less tractable than the reference species and lack well established genetic toolkits. The elevated nuclease activity observed here may represent a potential barrier to exogenous

### 3.2 Time-dependent exogenous DNA degradation in *Chlorella sorokiniana*

To investigate how rapidly exogenous DNA is degraded, plasmid DNA was incubated with cell lysates from *C. sorokiniana* and *P. tricornutum* over a time-course. As extensive DNA degradation was observed after 24 h in *C. sorokiniana* (Figure 1), shorter incubation times (5, 15, and 30 min, and 1 and 20 h) were examined to determine how quickly DNA degradation occurs under the assay conditions. *P. tricornutum* lysates were analysed after 1, 24, 48, and 96 h to determine whether DNA degradation occurred over an extended incubation period.

As shown in Figure 2B, *C. sorokiniana* lysates exhibited rapid degradation of plasmid DNA. After 5 min of incubation, only two faint plasmid DNA bands remained visible relative to the no-lysate control, indicating that substantial DNA degradation had already occurred. By 15 min, the intact plasmid band DNA stability and could contribute to the persistent transformation challenges encountered in these species. was no longer detectable and was replaced by a diffuse smear that progressively shifted towards lower molecular-weight DNA fragments with increasing incubation time, consistent with continued DNA fragmentation. In contrast, *P. tricornutum* showed little to no detectable degradation of plasmid DNA even after 96 h, highlighting a marked difference in nuclease activity towards double-stranded DNA between the two species (Figure 2A).

**Figure 2.**
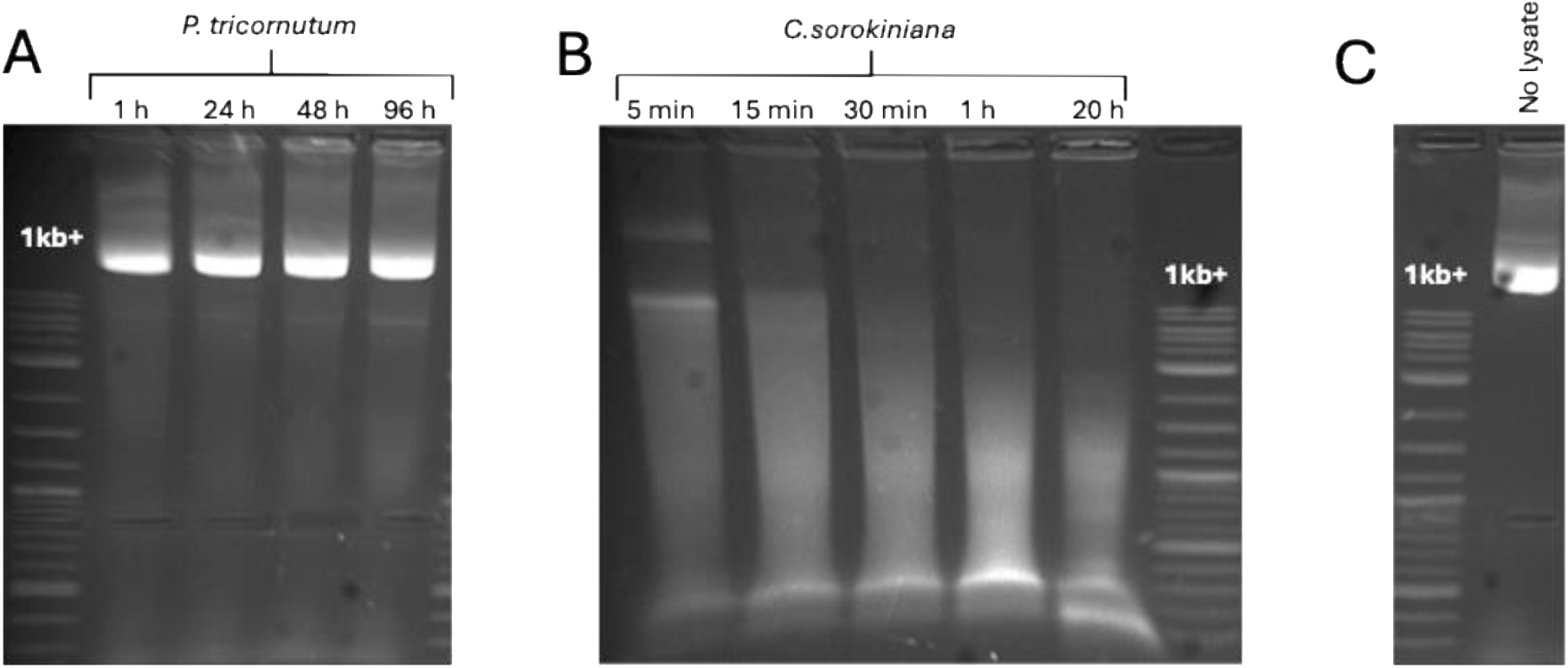
Time-course analysis of exogenous DNA degradation by *C. sorokiniana* and *P. tricornutum* cell lysates. (A) Plasmid DNA incubated with *P. tricornutum* lysates for 1, 24, 48, and 96 h. (B) Plasmid DNA incubated with *C. sorokiniana* lysates for 5, 15, and 30 min, and 1 and 20 h. (C) No-lysate controls for plasmid DNA. Equal amounts of total protein were used in each lysate reaction. Individual gels are labelled according to the algal species, and each lane is labelled with the corresponding incubation time. DNA size markers correspond to the 1 kb Plus DNA Ladder (New England Biolabs).

These findings demonstrate that *C. sorokiniana* lysates contain nuclease activity capable of rapidly degrading double-stranded DNA in vitro. However, as this assay uses whole-cell lysates, the observed activity may not fully reflect the in vivo situation, where other cellular factors could modulate nuclease access to exogenous DNA. An additional consideration is that freshwater species were grown in acetate-containing TAP under mixotrophic conditions, whereas marine species were grown phototrophically in ASW medium ^26^. These differences in growth conditions may influence cellular gene expression and nuclease abundance. Nonetheless, the rapid degradation of double-stranded DNA in *C. sorokiniana*, its absence in the three transformable reference species, and the intermediate activity observed in the recalcitrant *T. obliquus* strains suggest an association between nuclease activity and transformation recalcitrance (Figure 1). This

### 3.3 Inhibition of DNA-degrading activity by EDTA and Zn^2+^

To gain insight into the nature of the nucleases responsible for exogenous DNA degradation, plasmid DNA was incubated with *C. sorokiniana* and *T. obliquus* TO/164 cell lysates in the presence of EDTA (10 mM) or Zn^2+^(5 mM). Plasmid DNA incubated without lysate served as a negative control, and DNA degradation was assessed by agarose gel electrophoresis following 24 h incubation.

As shown in Figure 3, extensive plasmid DNA degradation was observed in lysates lacking inhibitors. In contrast, the presence of either EDTA or Zn^2+^ markedly reduced or completely inhibited DNA degradation, with intact plasmid bands observed after 24 h in both *C. sorokiniana* and *T. obliquus* (TO/164) lysates. EDTA and Zn^2+^ inhibit nuclease activity by mechanistically distinct routes — EDTA pattern supports nuclease-mediated DNA degradation as a potential barrier to the stability of exogenous DNA during transformation. chelates the divalent cations required for catalysis, whereas Zn^2+^ competes with the catalytic metal ions at the active site ^27–30^. The convergence of both inhibitors on the same activity therefore provides independent evidence that the observed nuclease activity is metal ion-dependent.

**Figure 3.**
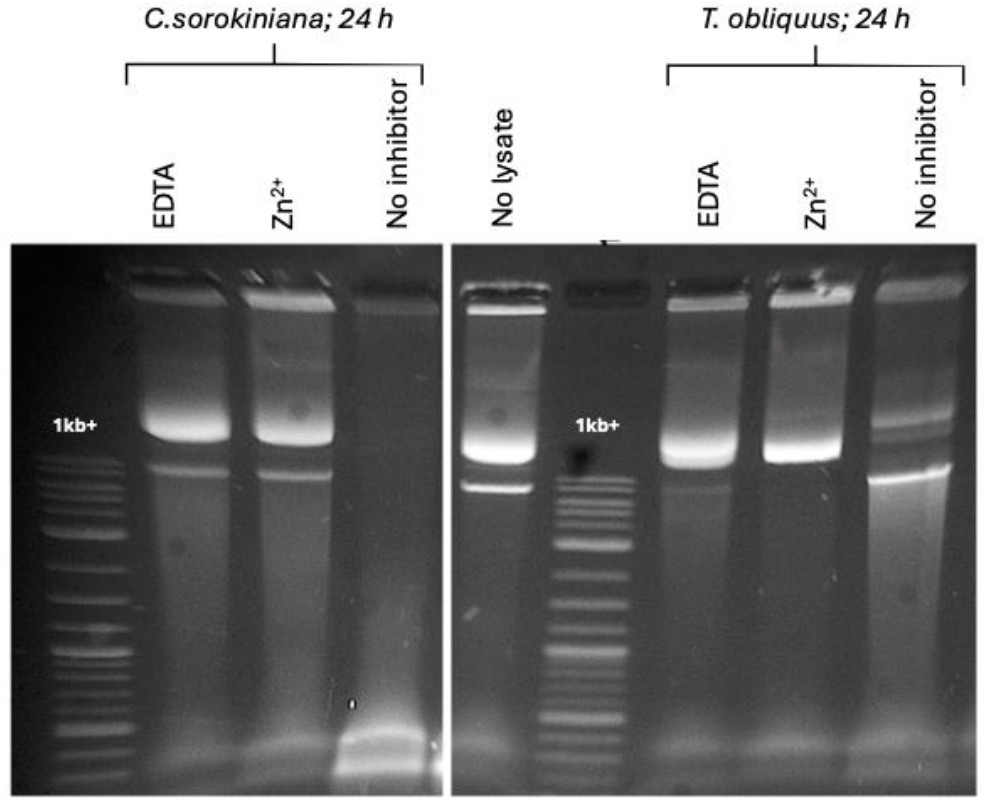
Effect of EDTA and Zn^2+^ on plasmid DNA degradation by *Chlorella sorokiniana* and *Tetradesmus obliquus* TO/164 cell lysates. Plasmid DNA was incubated with cell lysates in the absence or presence of EDTA or Zn^2+^ for 24 h. Equal amounts of plasmid DNA and total protein were used in each reaction. Individual gels are labelled according to the algal species, and each lane is labelled with the corresponding inhibitor treatment. No-lysate controls were included as negative controls. DNA size markers correspond to the 1 kb Plus DNA Ladder (New England Biolabs).

Together with the DNA fragmentation pattern observed in the time-course experiments (Fig. 2), the metal ion dependence and Zn^2+^ sensitivity is consistent with the involvement of a Mg^2+^/Ca^2+^-dependent endonuclease, such as a DNase I-like enzyme ^29^. The similar inhibition profiles observed in *C. sorokiniana* and *T. obliquus* TO/164 suggest that a similar class of nuclease may contribute to dsDNA degradation in both transformation-recalcitrant species. However, further biochemical and genetic analyses and identification of the responsible gene(s), will be required to determine the identity of the nuclease(s).

## 4 Conclusions

Collectively, these findings demonstrate that intracellular nuclease activities vary substantially among algal species and are greatest in transformation-recalcitrant species. *C. sorokiniana* exhibited rapid degradation of exogenous double-stranded DNA, while both *C. sorokiniana* and *T. obliquus* displayed a common metal ion-dependent nuclease activity consistent with a Mg^2+^/Ca^2+^-dependent endonuclease. Although multiple factors are likely to influence transformation efficiency, these findings suggest that intracellular nuclease activity may represent a barrier to exogenous DNA stability during transformation. However, the extent to which nuclease activity observed in whole-cell lysates reflect in vivo access to delivered DNA remains to be determined. Strategies aimed at improving intracellular DNA persistence, such as transient nuclease inhibition, DNA modification and/or protection, optimisation of growth conditions, or targeted disruption of nuclease genes, may provide potential approaches for enhancing transformation efficiency in genetically challenging algal species. Conversely, the rapid dsDNA-degrading activity observed in thermotolerant *C. sorokiniana* may have biotechnological value as a potential nuclease source if the responsible enzyme can be identified and further characterised.

## Acknowledgements

This work was funded by Innovate UK (Innovate UK Project number: 10160922)

